# The N400 ERP component reflects a learning signal during language comprehension

**DOI:** 10.1101/2021.03.25.436922

**Authors:** Alice Hodapp, Milena Rabovsky

**Author notes:** **Corresponding author:** Alice Hodapp, Department of Psychology, University of Potsdam, 14476 Potsdam, Germany.

## Abstract

The functional significance of the N400 ERP component is still actively debated. Based on neural network modeling it was recently proposed that the N400 component can be interpreted as the change in a probabilistic representation corresponding to an internal temporal-difference prediction error at the level of meaning that drives adaptation in language processing. These computational modeling results imply that increased N400 amplitudes should correspond to greater adaptation. To investigate this model derived hypothesis, the current study manipulated expectancy in a sentence reading task, which influenced N400 amplitudes, and critically also later implicit memory for the manipulated word: reaction times in a perceptual identification task were significantly faster for previously unexpected words. Additionally, it could be demonstrated that this adaptation seems to specifically depend on the process underlying N400 amplitudes, as participants with larger N400 differences also exhibited a larger implicit memory benefit. These findings support the interpretation of the N400 as an implicit learning signal in language processing.

## 1. Introduction

Neural computations have been suggested to be generally predictive (Friston, 2005). When the input deviates from probabilistic expectations, prediction errors occur, which are often thought of as graded rather than all-or-nothing phenomena and allow for an adaptation of the current model. This error-driven implicit learning could be the basis of our ability to adjust to the statistical regularities in our environment and ever-changing input. Prediction errors driving adaptation are suggested to be reflected in various event-related brain potentials (ERPs) in human electroencephalography (e.g., feedback related negativity: Chase et al., 2010; Cohen & Ranganath, 2007; Walsh & Anderson, 2012; error related negativity: Holroyd & Coles, 2002; mismatch negativity: Garrido et al., 2009; Wacongne et al., 2012). In the domain of language comprehension, the N400 ERP component has recently been interpreted as an implicit semantic prediction error driving adaptation of future expectations (Rabovsky et al., 2018; Rabovsky & McRae, 2014).

The N400 is a centro-parietally distributed negative-going ERP component that peaks around 400 ms after stimulus presentation and is modulated by a word’s predictability within a given context, with larger (more negative) amplitudes for contextually anomalous than for predictable words (Kutas & Hillyard, 1980). Despite the large number of studies on the N400, its functional significance is still actively debated (for a review see Kutas & Federmeier, 2011). Here, we focus on predictions derived from a neural network model of sentence comprehension, the Sentence Gestalt model (St. John & McClelland, 1990), which was used to model N400 amplitudes as the update in a probabilistic representation of sentence meaning implemented as the magnitude of change in the model’s hidden layer activation. The change in activation induced by a word corresponds to an implicit internal temporal difference prediction error at the level of meaning that drives adaptation of the model’s connection weights (Rabovsky et al., 2018). It is important to note that implicit prediction error is used here in the sense of Bayesian surprise (Itti & Baldi, 2009), i.e., the amount of unpredicted semantic information, and not in the sense of an expectancy violation per se (see Kuperberg & Jäger, 2016; Rabovsky et al., 2018; Rabovsky & McClelland, 2020; Rabovsky & McRae, 2014, for discussion). The N400 is thus simulated as the learning signal driving adaptation of our internal model of probabilities in the world to make better predictions in the future. Therefore, the model predicts that larger N400 amplitudes lead to greater adaptation and implicit learning.

One way to test this model derived prediction is to make use of the N400’s sensitivity to expectancy manipulations. The well-established finding of an interaction between expectancy and repetition can offer some insights into a possible relationship between N400 amplitudes and implicit learning. In these studies, incongruent sentences lead to a larger N400 amplitude at first presentation compared to a congruent sentence but also to a larger reduction in N400 amplitude when the sentence is repeated after a delay (Besson et al., 1992). A similar pattern emerges when the critical word is repeated while embedded in a different contextually unconstrained sentence (Rommers & Federmeier, 2018a). Interestingly, when unexpected words in a highly constrained context elicit similar N400 amplitudes as words in a weakly constrained sentence (not violating expectations), they also do not differ in magnitude of the repetition effect (Lai et al., 2021). This finding is in line with the view that it is not the expectation violation per se but rather the process underlying N400 amplitudes that is critical for model adaptation and implicit memory. While there are other possible explanations for this effect (Grill-Spector et al., 2006), the expectancy and repetition interaction was successfully simulated in the Sentence Gestalt model by Rabovsky et al. (2018), where the signal that was back-propagated through the network to drive learning corresponded exactly to the model’s N400 correlate, providing a computational explanation that needs further experimental investigation. The present study experimentally manipulates N400 amplitudes by varying expectancy, to directly test the hypothesis that the N400 reflects a learning signal in a subsequent behavioral implicit memory task.

## 2. Material and Methods

### 2.1. Participants

The study was preregistered on the Open Science Framework (https://osf.io/wg8nt/) and deviations are stated as such. The experiment is part of a research project whose protocols were approved by the Ethics Committee of the German Psychological Association (DGPs; MR102018). All participants gave written informed consent before the experimental session in accordance with the Code of Ethics of the World Medical Association (Declaration of Helsinki). 33 naive volunteers^1^ (12 male) participated in the experiment. Participants were compensated by either course credits or money (15€/h). Age ranged from 18 to 35 years (M = 23.88; SD = 5.60). All were native speakers of German and right-handed (as assessed via the Edinburgh Handedness Inventory). All had normal or corrected-to-normal vision and none reported a history of neurological or psychiatric disorders.

### 2.2. Stimuli and procedure

The stimuli consisted of 120 German sentences (See Table 1 for example sentences, Appendix A1 for the controlled stimulus characteristics, and the OSF for the complete list of sentences: https://osf.io/wg8nt/) that were created in pairs that ended with the same target noun. Nouns with the highest cloze probability in the unexpected sentences were never used as a target word in another sentence, as to prevent downstream consequences of expected but not seen words (Hubbard et al., 2019; Rommers & Federmeier, 2018b). The sentence pairs were divided across three counterbalanced lists, so that each target word was presented to an individual participant in either its expected or unexpected version, or not presented at all (serving as a not seen control condition for the implicit memory task later). Each of the lists contained 40 expected, 40 unexpected and 50 filler sentences. Sentences were divided into part A (first half of the sentence list) and part B (second half), which contained half of each stimulus condition respectively. The order of part A and B was counterbalanced across participants and was kept the same for the implicit memory task. Sentence order was randomized within participants and it was additionally ensured that no more than two unexpected sentences would be presented successively.

**Table 1.** Example sentences and their approximate translations into English. The expected or unexpected target words are underlined.

|  |  |
| --- | --- |
| expected | 1 a. Der Zauberer holte ein Kaninchen aus seinem <u>Hut</u><br>*The magician took a rabbit out of his <u>hat</u>* |
| unexpected | 1 b. Der Tierwärter holte ein Kaninchen aus seinem <u>Hut</u><br>*The zookeeper took a rabbit out of his <u>hat</u>* |
| expected | 2 a. In Bayern angekommen, aß der Tourist eine frisch gebackene <u>Brezel</u><br>*After arriving in Bavaria the tourist ate a freshly baked <u>pretzel</u>* |
| unexpected | 2 b. In Italien angekommen, aß der Tourist eine frisch gebackene <u>Brezel</u><br>*After arriving in Italy the tourist ate a freshly baked <u>pretzel</u>* |
| expected | 3 a. Elias wollte Mia eine Postkarte schreiben und fragte nach ihrer <u>Adresse</u><br>*Elias wanted to write to Mia and asked for her <u>address</u>* |
| unexpected | 3 b. Elias wollte Mia anrufen und fragte nach ihrer <u>Adresse</u><br>*Elias wanted to call Mia and asked for her <u>address</u>* |
| expected | 4 a. Nach einer langen Regenzeit versprach der Wetterbericht nun endlich wieder <u>Sonne</u><br>*After a long period of rain, the forecast finally called for <u>sun</u>* |
| unexpected | 4 b. Nach einer langen Dürrezeit versprach der Wetterbericht nun endlich wieder <u>Sonne</u><br>*After a long period of drought, the forecast finally called for <u>sun</u>* |

The experiment was divided into three tasks and all stimulation was controlled by running the Psychophysics Toolbox (Brainard, 1997; Kleiner et al., 2007) in MATLAB R2020a (MathWorks Inc. Natick, MA, USA). In the sentence reading task, participants were instructed to attentively read each sentence for comprehension. Each trial started with a fixation cross until the participant pressed a button, after which a 300 ms black screen was presented. Then each word appeared for 250 ms at the center of the screen with an interstimulus interval of 300 ms. Word presentation time was adjusted for words with more than 12 letters (20 ms per additional letter). The sentence final word appeared for 300 ms and was followed by an 800 ms interstimulus interval before the fixation cross reappeared. Participants were unaware of the upcoming implicit memory task while reading the sentences.

After the sentence task, participants completed 4 blocks of an n-back task with numbers to minimize any potential explicit memory effects of sentence reading. In the n-back task participants are instructed to respond when a stimulus matches the one that appeared n items before, which increases working memory load with each block (Kirchner, 1958).

Participants’ implicit memory for the target words was then assessed via a behavioral perceptual identification paradigm adapted from Stark & McClelland’s (2000) work on repetition priming. Perceptual identification tests are a reliable measure of implicit memory (Buchner & Wippich, 2000) and the adapted paradigm used here showed robust priming effects on reaction times (Stark & McClelland, 2000). Each participant was presented with all 120 target words that were used across lists. Again, the order of the stimuli was randomized within participant, however the overall order of part A and B stimuli (as compared to the sentence reading task) stayed the same, to control for strong timing effects. Participants started each trial with a button press. After a 300 ms interval with a blank screen, a mask (15 #s, two more than the longest target word) was presented for 14 frames immediately followed by the first target word presentation for one frame (1 frame = 16.67 ms). This procedure continued, until the participants pressed a button to indicate that they had recognized the word, but with each round the mask duration was decreased by one frame, whereas the stimulus presentation was increased by one (see illustration in Figure 1). This allowed the recording of reaction times corresponding to a word’s perceptual fluency. Participants were instructed to identify the word as early on and as accurately as possible. After button press, a cue was presented on the screen, prompting participants to type in their response. Participants received feedback for incorrect and too slow trials (after 25 repetitions the word became fully visible and the trial aborted) only.

**Figure 1.**
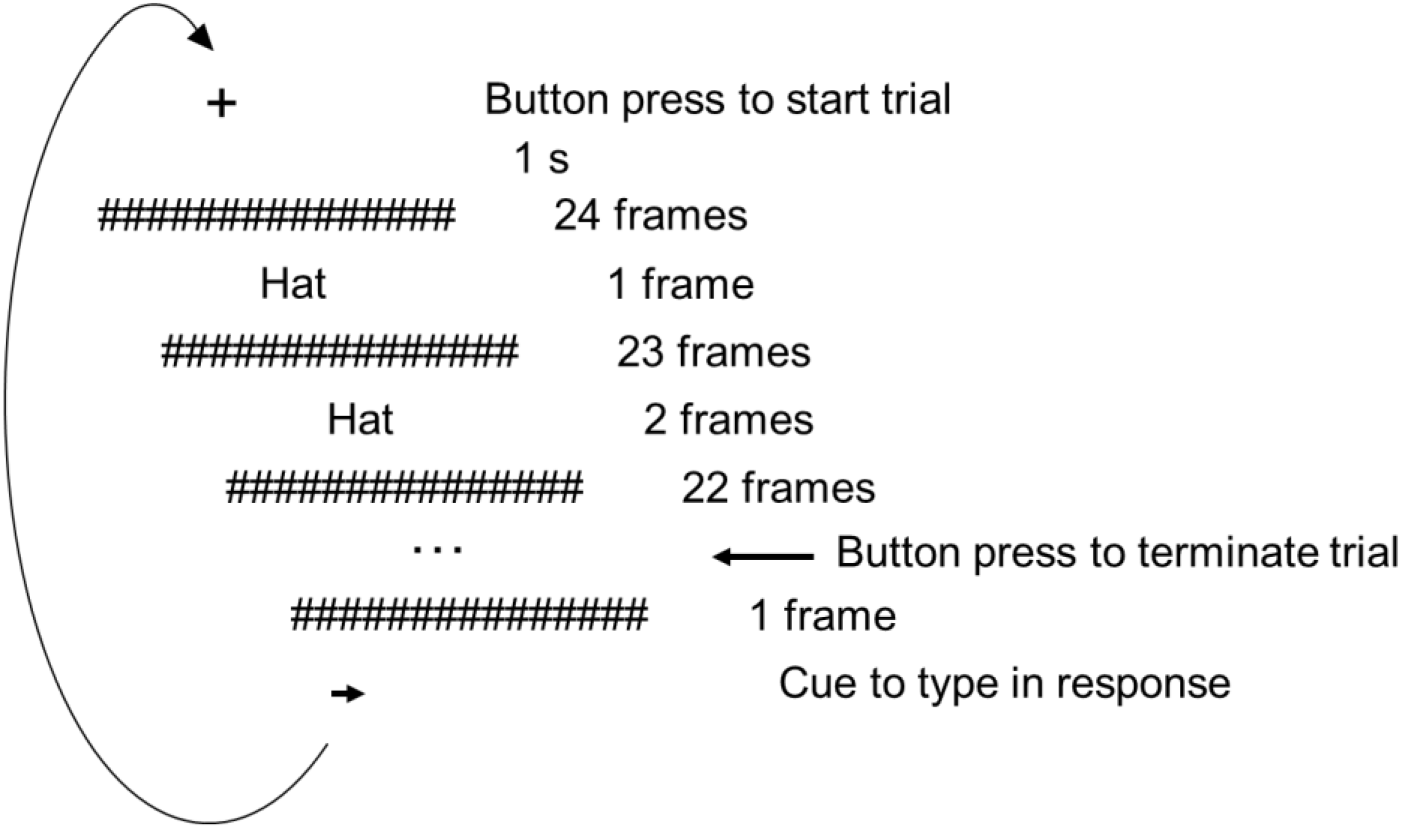
Design of the perceptual identification task as adapted from Stark & McClelland (2000). The word becomes increasingly clear with time until the participant presses a button. They are then instructed to type in the word that they had recognized. The design allows the recording of continuous reaction time data which has been shown to be modulated by implicit memory.

### 2.3. Data acquisition and analysis

#### EEG recording

During sentence reading, participant’s electroencephalogram (EEG) was recorded continuously from 64 active Ag/AgCl electrodes positioned according to the extension of the international 10-20 system. The EEG was referenced online to the left mastoid and electrode impedances were kept below 5 kΩ. The data were acquired at a sampling rate of 1000 Hz and amplified (BrainAmp amplifier with a bandpass filter of .016-250 Hz and a time constant of 10s).

#### EEG analysis

EEG data were pre-processed and analyzed with MATLAB R2020a using the EEGlab (Delorme & Makeig, 2004) toolbox. Data were re-referenced offline to the average of the left and right mastoids. The EEG was filtered with a 0.1 Hz high-pass filter (two-pass Butterworth with a 12 dB/oct roll-off) and low-pass filtered at 30 Hz (two-pass Butterworth with a 24 dB/oct roll-off). The continuous EEG data were then segmented into epochs ranging from −200 to 1000 ms relative to target word onset. A 200 ms baseline was subtracted. Eye blinks were corrected by means of independent component analysis (Jung et al., 2000; Makeig, Jung, Gahremani, Bell, & Sejnowski, 1997). Bad channels were identified by their variance (in case it exceeded an absolute z-score of 3) and were interpolated using a spherical spline function (Perrin et al., 1989). All segments with values that exceeded ± 75 μV at any channel were excluded. N400 data were analyzed for a centro-parietal region of interest (ROI: CPz, Cz, CP1, CPz, CP2, P1, Pz, P2) within a 300 to 500 ms time window.

#### Statistics

We performed a linear mixed-effects model (LMM) analysis using the package lme4 as implemented in R (R core Team, 2018) to investigate the effect of experimental condition on N400 amplitude and implicit memory, as well as the overall correlation between N400 and implicit memory. Reaction times for correct responses were log-transformed (due to a skewed distribution; not preregistered). For all models predicting reaction times, word frequency was added into each model as additional fixed effect (log transformed, centered, and scaled). Following the recommendations of Barr et al. (2013), we tried to fit the maximal random effect structure as justified by the design but removed random correlations to aid convergence for the analysis of a condition effect on reaction times. Sum coding (−0.5, 0.5; for EEG analysis) and Helmert coding (for behavioral analysis) were used as contrasts. The significance of fixed effects was determined via likelihood ratio tests to compare the fit of the model to that of a model with the same random effects structure without the respective fixed effect.

## 3. Results

Figure 2 shows the ERP time-locked to the onset of the target word. In line with previous studies, a clear N400 is visible and modulated by expectancy condition. The amplitude of the N400 in response to unexpected words was more negative than for expected words by 1.32 μV (*SE* = .375, *t* = −3.511, *χ^2^* = 10.475, *p* = .001).

**Figure 2.**
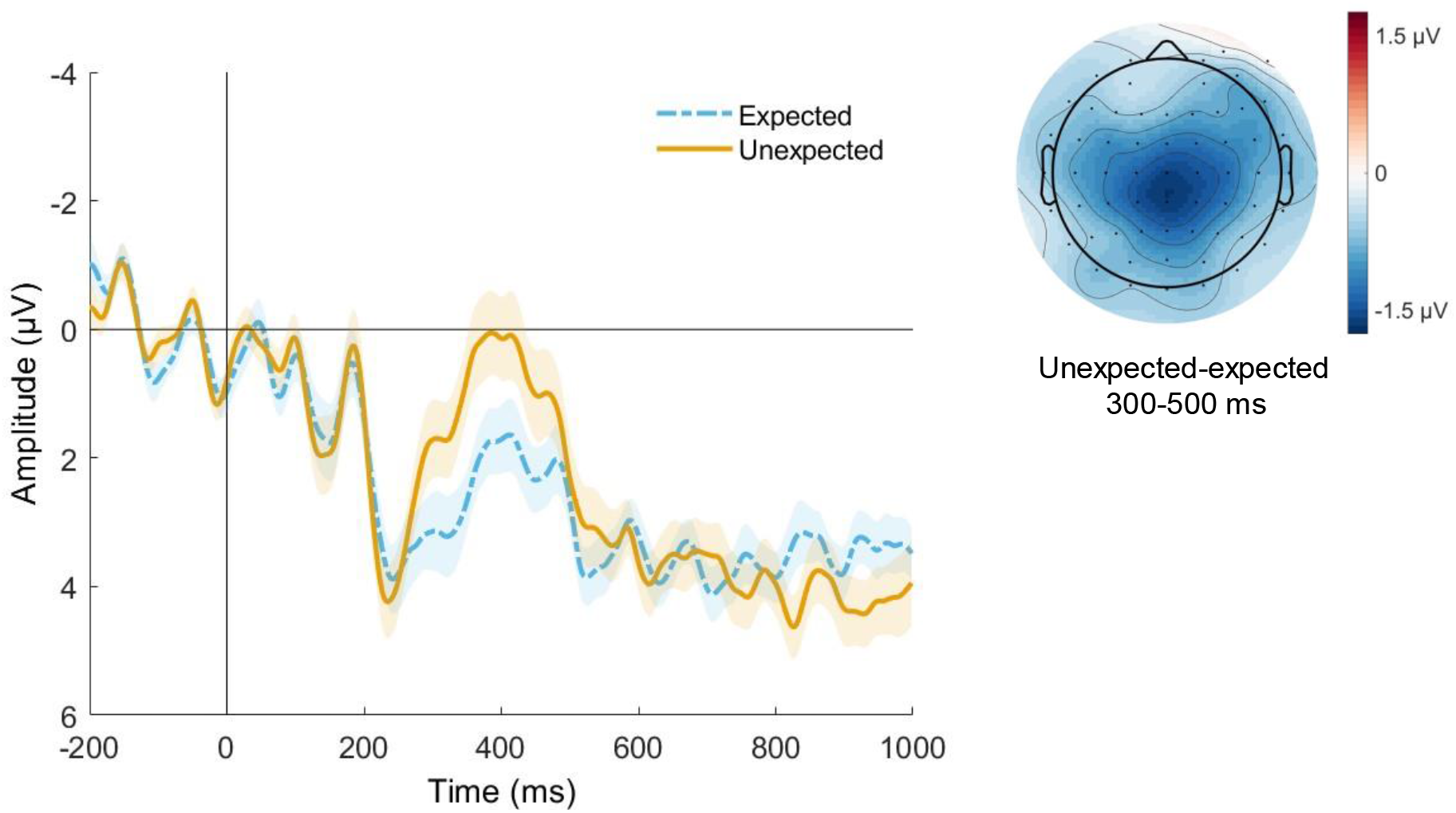
Grand-average waveforms (n = 33) at centro-parietal ROI and topography for the 300-500ms time window. Note that negative values are plotted upwards. Error bands indicate the SEM.

The arithmetic means of the reaction time data show a 96 ms benefit for previously unexpected compared to not seen and a 79 ms benefit for previously unexpected compared to expected words (Table 2). Log transformed data is depicted in Figure 3. In a LMM analysis on log-transformed data, the behavioral results showed the classic priming effect (not seen words vs. all previously seen words) on reaction times: *β* = − 0.024, *SE* = 0.008, *t* = −2.853, *χ^2^* = 7.244, *p* = .007^2^. Critically, previous expectancy of the primed words influenced reaction times: previously unexpected words were recognized faster than previously expected words (*β* = −0.037, *SE* = 0.008, *t* = −4.557, *χ^2^* = 17.764,*p* < .001). There was an additional main effect of word frequency (*β* = −0.026, *SE* = 0.008, *t* = −3.500, *χ^2^* = 11.540,*p* < .001). Error rates did not differ between conditions.

**Figure 3.**
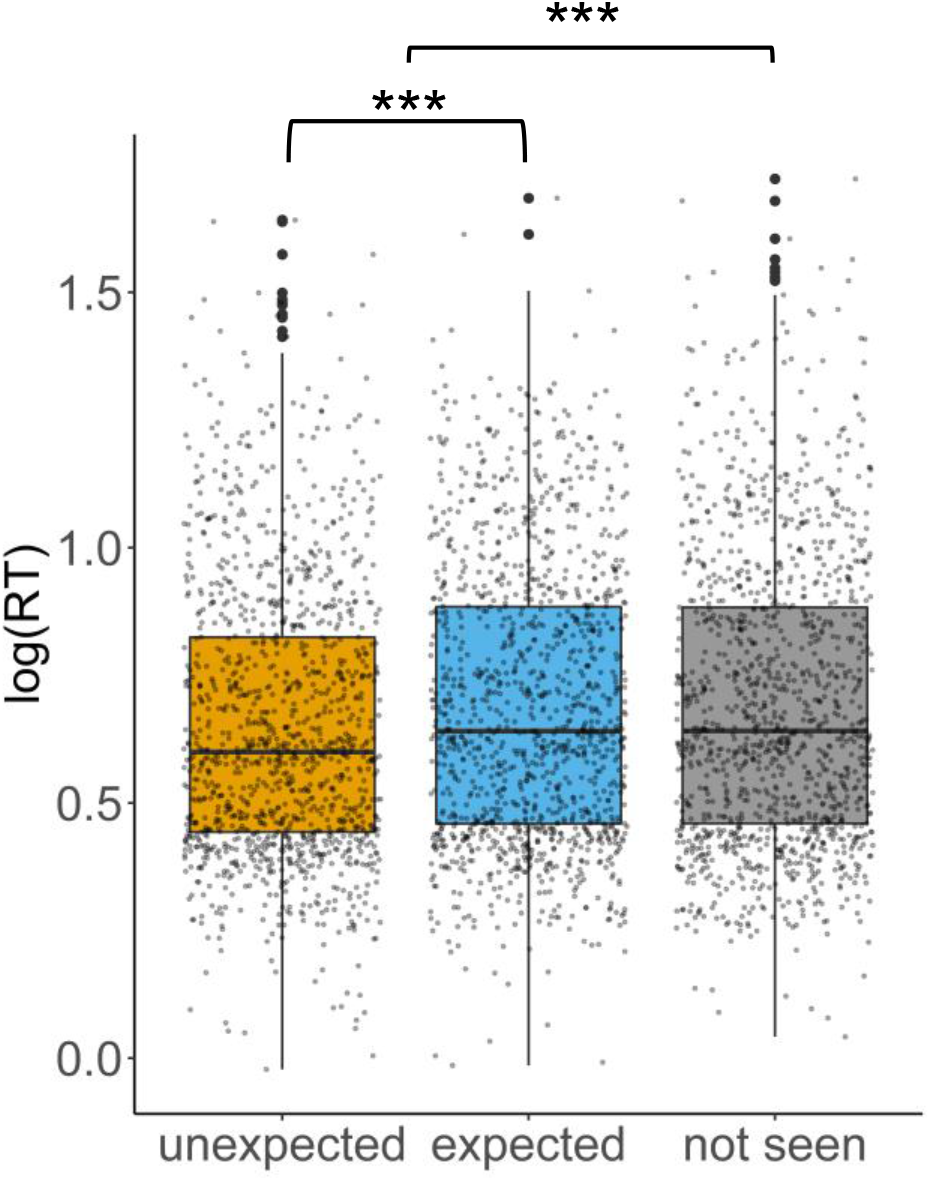
Reaction times (log transformed) in the perceptual identification task across participants and items by condition, i.e., unexpected, expected or not seen during the preceding sentence reading task.

**Table 2.** Mean reaction times and error rates in the perceptual identification task per condition, i.e., unexpected, expected or not seen during the preceding sentence reading task.

|  | Unexpected | Expected | Not seen |
| --- | --- | --- | --- |
| Reaction time in ms | 1990 (597) | 2069 (606) | 2086 (655) |
| Error rate in % | 4.7 (4.0%) | 4.2 (3.5%) | 4.9 (4.4%) |

There was no main effect of N400 amplitude during sentence reading on reaction times in the perceptual identification task (*β* = −0.0007, *SE* = 0.0005, *t* = −1.495, *χ^2^* = 2.019, *p* = .155)^3^ when controlling for the main effect of frequency (*β* = −0.021, *SE* = 0.006, *t* = −3.474, *χ^2^* = 11.068, *p* < .001)^4^. An additional exploratory analysis revealed a significant positive correlation between participants’ individual N400 amplitude differences (expected minus unexpected) during sentence reading and the respective difference in reaction times in the perceptual identification task with *r* = 0.46 [95 % CI: 0.14, 0.70], *p* = .007 (Figure 4A). No support for such a correlation was found for (frontal and parietal) post N400 positivities that have been reported for unexpected sentence continuations (Kuperberg et al., 2019; Van Petten & Luka, 2012). For the late frontal positivity results see Figure 4B (there was no significant parietal P600 effect in our data; for details on both components see Appendix A2).

**Figure 4.**
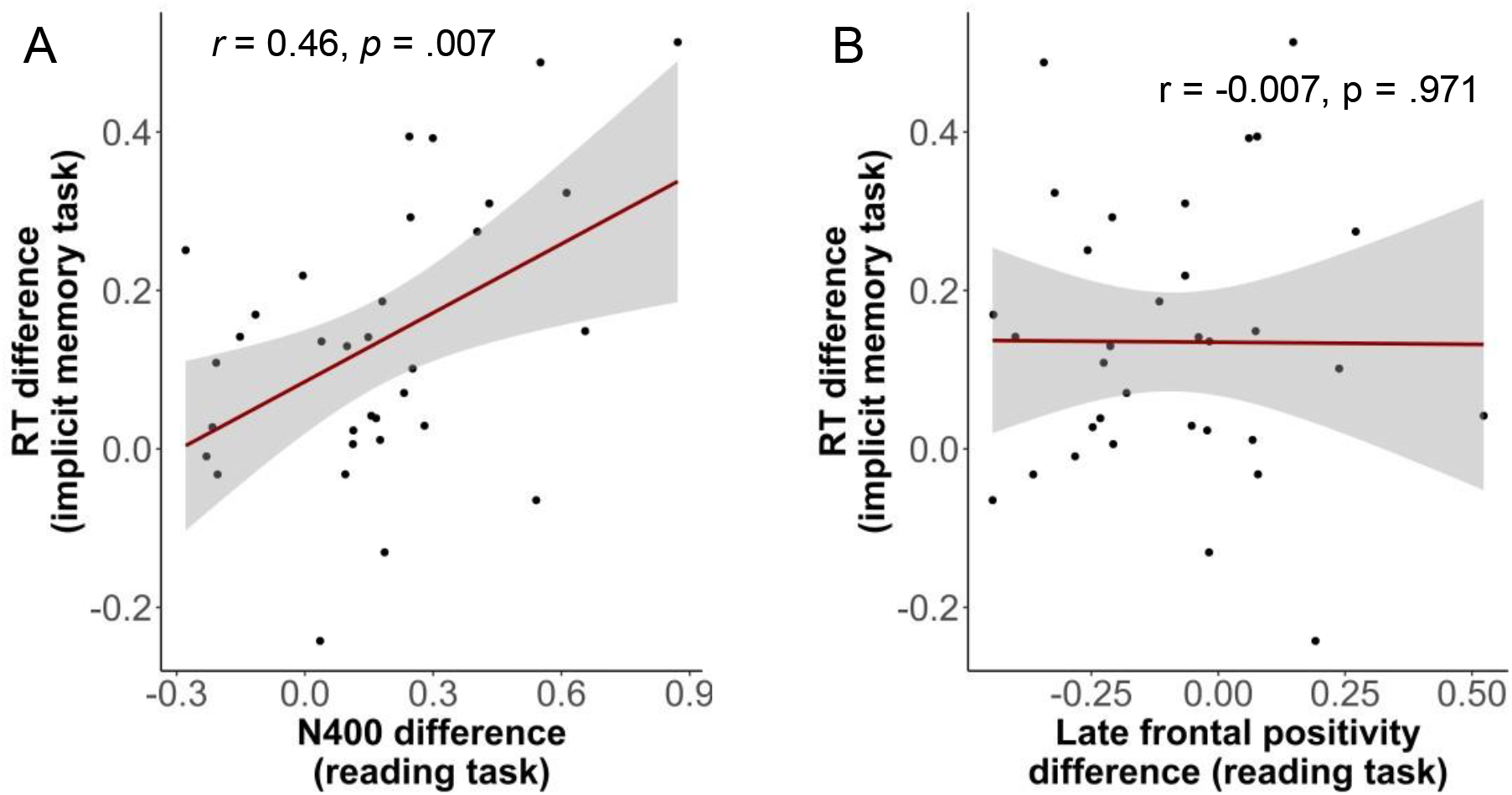
Correlation of within participant (A) N400 differences (B) late frontal positivity differences (both expected minus unexpected) in the reading task and the respective reaction time difference from the subsequent implicit memory task. Error bands indicate the SEM.

## 4. Discussion and conclusion

This study tested the hypothesis that N400 amplitudes reflect a learning signal by experimentally manipulating word expectancy during sentence reading and subsequently presenting the critical words in a perceptual identification task to measure implicit memory. In line with well-established findings in the literature on the N400, the experimental manipulation successfully influenced N400 amplitudes (Figure 2). As predicted, this manipulation also influenced participants’ implicit memory and thus previously unexpected words were recognized faster than expected words or words that had not been encountered before (Figure 3). The results are consistent with the assumption that the N400 reflects an implicit semantic prediction error that drives model adaptation (Rabovsky et al., 2018; Rabovsky & McRae, 2014) and unexpected words leading to larger N400 amplitudes and enhanced implicit memory.

This theory is also supported by results from studies that were not explicitly designed to investigate this relationship. In a study aiming to investigate ERP correlates of memory encoding, the authors observed a relation between a N400-like negativity for single words presented during the study phase and subsequent implicit memory performance as reflected in a stem completion task during test (Schott et al., 2002). Meyer et al. (2007) found a positive correlation between the amplitude of the N400 at encoding and the size of the early old/new ERP effect at test. Further support for the N400 as a learning signal comes from the field of language learning in early childhood: The presence of N400 effects as well as N400 priming effects seem to be predictive of infants’ later language skills (Friedrich & Friederici, 2006, 2010). Even though these results were not originally interpreted this way, they can be naturally explained by understanding the N400 as an implicit semantic prediction error that drives implicit learning.

The results suggest that word expectancy does not independently influence both N400 amplitudes and adaptation. The correlation between N400 amplitude differences and subsequent implicit memory (but not between post N400 positivity differences and implicit memory; Figure 4) implies that participants with overall larger N400 effects also exhibit larger adaption effects, supporting the theory that N400 amplitudes reflect the learning signal driving model adaptation.

In sum, the current study found experimental support for a prediction derived from computational modeling work (Rabovsky et al., 2018; Rabovsky & McRae, 2014), namely that the N400 ERP component reflects a learning signal during language comprehension.

## Data and Code Availability

Grand averaged EEG files, data, and scripts needed for statistical analysis can be found on
OSF (https://osf.io/wg8nt/). Due to the European Data Protection Law, raw data is only available from the corresponding author upon request.

## Acknowledgments

Funding was provided by an Emmy Noether grant from the German Research Foundation (grant RA 2715/2-1) to Milena Rabovsky. We thank Antonia Heinrich for help with stimulus preparation.

## Footnotes

1 We preregistered 42 participants, but due to the COVID-19 pandemic data acquisition had to stop and we decided to analyze the data of the 33 participants for which we obtained data prior to the shutdown.

2 For completeness: unexpected words (*β* = −0.042, *SE* = 0.008, *t* = −5.181, *χ2* = 22.51, *p* < .001) but not previously expected words (*β* = −0.005, *SE* = 0.008, *t* = −0.636, *χ*2 = 0.403, *p* = .526) were recognized significantly faster than not seen words.

3 The missing correlation between N400 amplitudes and reaction times can be explained by the intuitive notion that words which are easier to access in semantic memory elicit a lower amplitude N400 (Kutas & Federmeier, 2011; Van Berkum, 2009) and are also easier to identify in e.g., a perceptual identification task.

4 The slight deviance in frequency effect (with respect to the frequency effect in the main analysis of the behavioral results) is due to the analysis only containing trials which survived artifact rejection in EEG-pre-processing.

## Notes

### Competing Interest Statement

The authors have declared no competing interest.

