## Appendix A1 for "The N400 ERP component reflects a learning signal during language comprehension"

Table A.1. Stimuli characteristics for the expected and unexpected condition and for the three stimulus lists.

| | List 1 | List 2 | List 3 | Expected | Unexpected |
|---|---|---|---|---|---|
| Cloze probability | 0.41 (0.43) | 0.40 (0.40) | 0.39 (0.40) | 0.79 (0.18) | 0.01 (0.04) |
| Plausibility | 4.47 (2.15) | 4.43 (2.17) | 4.67 (1.97) | 6.39 (0.61) | 2.66 (1.18) |
| Sentence length | 12.56 (2.93) | 12.98 (2.67) | 12.19 (2.52) | 12.54 (2.71) | 12.60 (2.74) |
| Frequency normalized | 30.78 (91.73) | 21.37 (32.75) | 34.78 (93.09) | 29.00 (77.83) | |
| Neighborhood size | 12.7 (15.4) | 13.6 (15.9) | 15.4 (17.7) | 14.13 (16.35) | |
| Target length | 7.02 (2.59) | 7.25 (6.22) | 6.61 (2.30) | 7.18 (5.32) | |

Notes. Since all sentences were created in pairs, all lexical measures, semantic features, and visual input for the target word were the same for both conditions. Cloze probability was operationalized as the proportion of participants (n=25) who used the critical word as sentence continuation in an online cloze test (on www.prolific.co). Annotated type frequency (normalized) and orthographic neighborhood size (as defined by Coltheart et al., 1977) according to dlexDB (www.dlexdb.de).
