## Appendix A2 for "The N400 ERP component reflects a learning signal during language comprehension"

Exploratory (not preregistered) analysis of post N400 positivities related to unexpected sentence continuations (Kuperberg et al., 2019; Van Petten & Luka, 2012).

*Late frontal positivity*. A possible frontal post N400 positivity was analyzed in a frontal ROI (FP1, FP2, AF3, AF4). There was a significant effect of expectancy (Figure A1; expected vs. unexpected: $\beta = 0.790$, $SE = 0.324$, $t = 2.440$, $\chi^2 = 5.715$, $p = .017$) in a 600-1000 ms time window but no significant correlation between within participant differences in the late frontal positivity and reaction time differences in the implicit memory task (Figure 4B; $r = -0.007$ [95 % CI: -0.35, 0.34], $p = .971$).

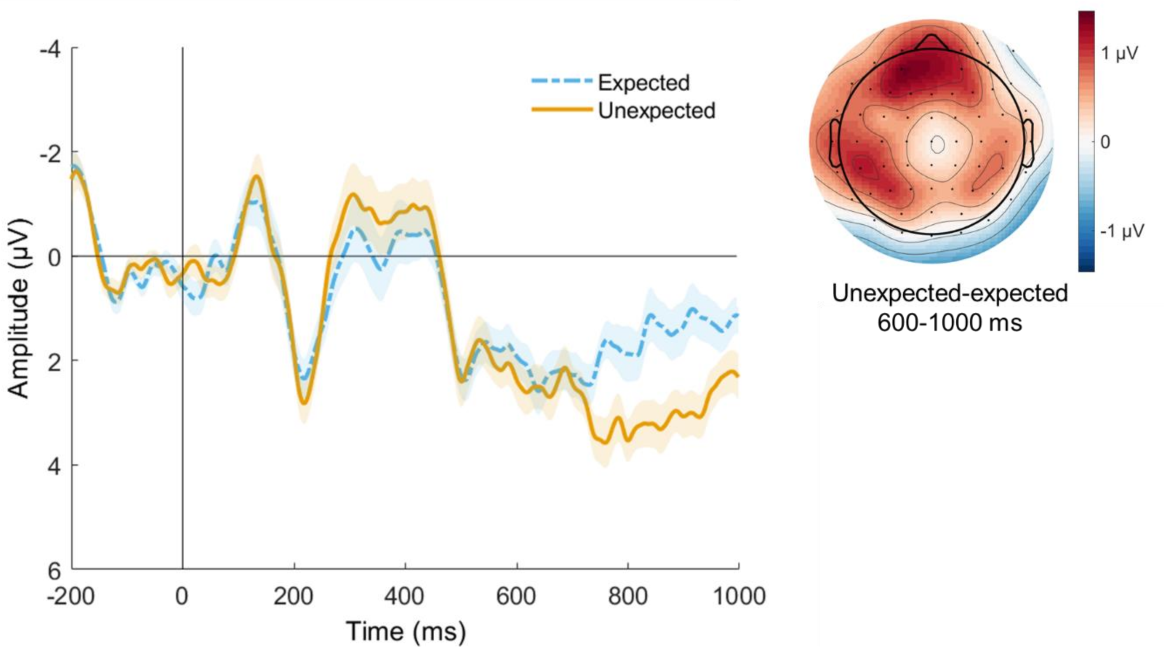


Figure A1. Grand-average waveforms (n=33) at frontal ROI and topography for the 600-1000 ms time window. Note that negative values are plotted upwards. Error bands indicate the SEM.

*P600 (late parietal positivity)*. There was no significant P600 effect (expected vs. unexpected: $\beta = 0.267$, $SE = 0.288$, $t = 0.926$, $\chi^2 = 0.855$, $p = .355$) in a parietal ROI (P3, P1,

P2, Pz, P4, PO3, POz, PO4, O1, O2, Oz) in a literature-based time window of 600-800 ms (Figure A2). There was no significant correlation between within participant differences in the late parietal positivity and reaction time differences in the implicit memory task (*r* = 0.32 [95 % CI: -0.03, 0.60], *p* = .072). Note that the trend in correlation is opposite of what would be expected if there were an effect of this parietal positivity on implicit memory in the sense that a larger late parietal positivity for unexpected words would entail a stronger implicit memory benefit for these words. Instead, the positive trend is likely driven by participants with rather late or widespread N400 effects.

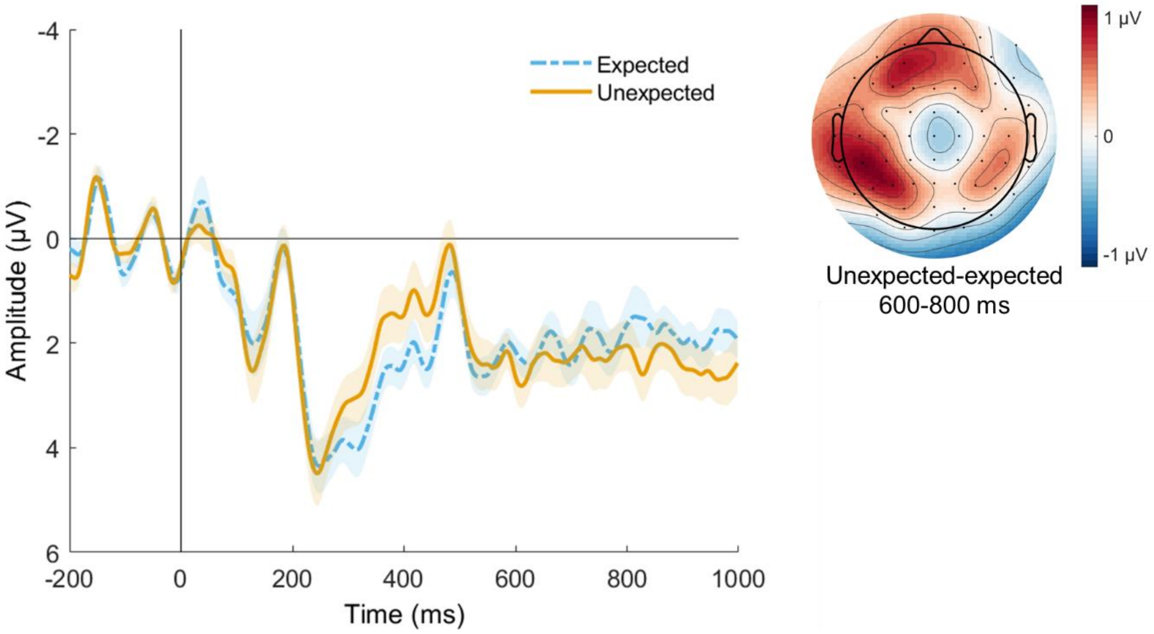


Figure A2. Grand-average waveforms (n=33) at parietal ROI and topography for the 600-800 ms time window. Note that negative values are plotted upwards. Error bands indicate the SEM.
